# Beyond size dimorphism: the past, present, and future of Rensch’s Rule

**DOI:** 10.1101/2024.10.30.621038

**Authors:** Ken S. Toyama

## Abstract

‘Rensch’s Rule’ is known as a pattern of allometry in which the degree of male-biased sexual size dimorphism (SSD) increases with species body size. Over the last decades, a growing amount of Rensch’s Rule studies has advanced our understanding of SSD, its prevalence in nature, and the mechanisms underlying its evolution. However, Bernhard Rensch, when describing the pattern for the first time, considered the allometry of SSD only as a special case of a more general pattern in which dimorphism in any relative sexual difference increased with body size. In this perspective I revisit the history of Rensch’s Rule, starting with its popularization in recent decades, then diving into the original works by Rensch to rediscover his original observations, and finally discussing the implications of studying Rensch’s pattern beyond its applications to SSD. The strong bias towards body size in the study of Rensch’s Rule has proven valuable regarding our understanding of the evolution of SSD. Using empirical examples I propose, however, that expanding the study of the pattern to other traits might prove insightful for the general study of sexual dimorphism and phenotypic diversity.

## Introduction

‘Rensch’s Rule’ is the name given to an interspecific pattern of allometry in which the degree of male-biased sexual size dimorphism (SSD) increases with species size (Abouheif and Fairbairn, 1997, Fairbairn 1997, Meiri and Liang, 2021; Figure 1). The study of Rensch’s Rule has become particularly popular in the last three decades, as hundreds of studies have tested the existence of the pattern in different taxa, as well as possible hypotheses for its emergence. Rensch’s Rule was once believed to be a generalizable pattern among animals (Abouheif and Fairbairn, 1997). Examples of taxa in which Rensch’s Rule has been tested and confirmed include some groups of primates, lizards, birds, and insects (e.g. Abouheif and Fairbairn, 1997; Smith and Cheverud, 2002; Blanckenhorn et al., 2007; Dale et al., 2007; Liang et al., 2022). However, other taxa, including amphibians and some groups of birds and lizards, do not show the pattern or even reverse it (e.g. Webb and Freckleton, 2007; De Lisle and Rowe, 2013; Liang et al., 2022). Regardless, the increasing interest in the allometric patterns of SSD found in nature has simultaneously stimulated the study of the processes underlying them and, consequently, has provided numerous insights on the evolution of size differences between the sexes.

**Figure 1.**
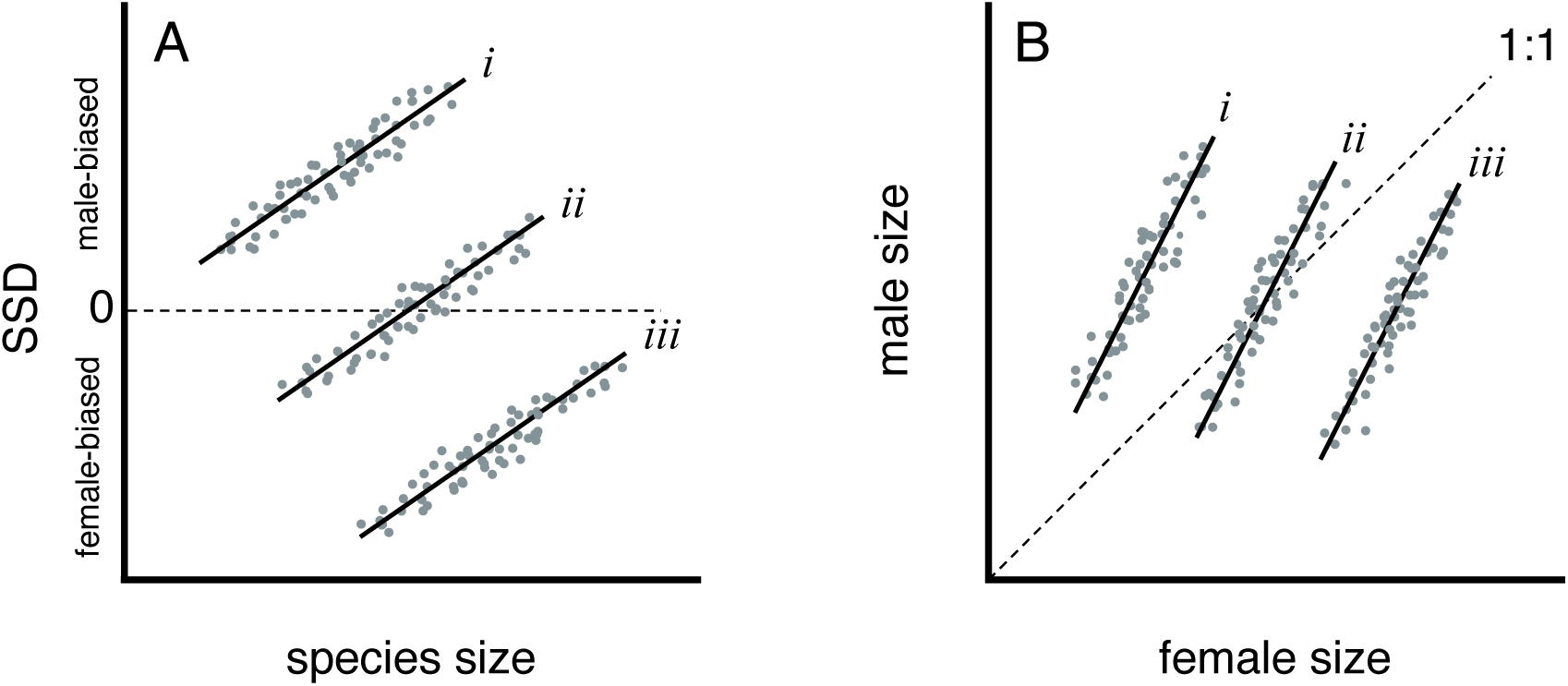
Three hypothetical clades in which body size follows Rensch’s Rule. (A) Rensch’s Rule is known as an interspecific pattern of allometry in which male-biased sexual size dimorphism (SSD) increases with species size. The occurrence of the pattern is independent of whether species in a particular clade show (*i*) male-biased, (*ii*) mixed, or (*iii*) female-biased SSD. (B) Another way to illustrate Rensch’s Rule is by testing the relationship between log-transformed male and female body sizes. If female size is represented on the *x* axis, clades for which this relationship has a slope > 1 (where 1 represents isometry) are following Rensch’s Rule.

Rensch’s Rule is based on the observations of Bernhard Rensch, a German biologist famous for his work on diverse topics including macroevolution, biogeography, and biophilosophy, but especially for his contributions to the Modern Evolutionary Synthesis (Wuketits, 2006). Rensch’s observations regarding the rule that bears his name appeared first in two of his works: the article *Die Abhängigkeit der relativen Sexualdifferenz von der Körpergrösse* (Rensch, 1950; in German) and the second edition of the book *Neuere Probleme der Abstammungslehre. Die transspezifische Evolution* (Rensch, 1954; in German), the latter more widely known as its translated version: *Evolution Above the Species Level* (Rensch, 1959), and possibly his most influential work. These works suggest that Rensch was originally interested not only in the allometry of SSD, but in the allometry of any relative sexual difference (e.g., dimorphism in secondary sexual characters like horns, antlers, teeth, etc). In fact, he described how such relative sexual differences tended to increase in magnitude with absolute size in many taxa, proposing this pattern as an emerging macroevolutionary rule. Rensch treated dimorphism in body size just as another case of this general pattern; however, Rensch’s Rule eventually became known, almost exclusively, as an allometric pattern of SSD, overlooking the generality of Rensch’s original observations.

In this perspective I revisit the history of this iconic evolutionary pattern, focusing on the fact that its modern definition is, at least, incomplete. I start in the present, describing the beginnings of the ‘modern’ Rensch’s Rule and how it has driven the last three decades of research on the allometry of sexual dimorphism. Then, I look into Rensch’s foundational works on the subject and revisit his original ideas, contrasting them against the modern definition of Rensch’s Rule. Finally, I use empirical examples to illustrate how overlooking the generality of Rensch’s original pattern might limit our understanding of the processes driving the evolution of sexual dimorphism, and how complementing the ‘modern’ Rensch’s Rule with Rensch’s more general observations can enrich the way we explore the emergence of sexual differences at the macroevolutionary level.

### I. Rensch’s Rule: the modern interpretation of Rensch’s observations

Although the relationship between SSD and body size had already been the focus of influential research in the decades that followed the publication of Rensch’s seminal works (e.g., Clutton-Brock et al., 1977; Leutenegger, 1978; Fitch, 1981), the term ‘Rensch’s Rule’ was not popularized until the 1990s through a series of papers authored by Daphne Fairbairn and Ehab Abouheif. Fairbairn had already published on the allometry of SSD in the early 1990s (e.g. Fairbairn, 1990; Fairbairn and Shine, 1993; Fairbairn and Preziosi, 1994) but the first explicit mention of the term ‘Rensch’s Rule’ in this decade appeared in Abouheif’s Master’s thesis (1995), which was supervised by Fairbairn (notice, however, that the term ‘Rensch’s Rule’ had already been used as early as 1970; see Earhart and Johnson, 1970). Two subsequent papers (Abouheif and Fairbairn, 1997; Fairbairn, 1997) would become landmark works on the study of the allometry of SSD, as they established the methodological and conceptual bases for the study of Rensch’s Rule in the following decades. These works also became highly influential because they reviewed the available evidence for Rensch’s Rule in nature, which partly supported its generality across taxa.

Citing Bernhard Rensch’s book *Evolution above the species level* (1959) as the original source, the influential works of Abouheif and Fairbairn defined Rensch’s Rule as a pattern in which SSD increases with body size in taxa where males are the larger sex, but decreases with body size where females are the larger sex (Abouheif and Fairbairn, 1997; Fairbairn, 1997; Figure 1). They also popularized what for the past decades has been considered the standard way to test Rensch’s Rule: the examination of the allometric slope between log-transformed male and log-transformed female size among related species (Figure 1B) (but notice that this technique had already been used before, e.g. Leutenegger, 1978). This method was proposed given that regressing SSD on body size would break the assumption of independence between the dependent and predictor variables (but see Adams et al., 2020). In their papers, Abouheif and Fairbairn also brought up the issue of non-independence due to evolutionary relatedness, noting that failure to account for phylogenetic relationships, which was not uncommon in the preceding years, can lead to erroneous conclusions when testing Rensch’s Rule.

After the initial definitions and methodological considerations, Abouheif and Fairbairn’s papers proceed to show empirically that Rensch’s Rule is highly prevalent in nature and more pronounced in clades with male-biased SSD, which is consistent with the idea of sexual selection being a major driver of this pattern. Importantly, Fairbairn (1997) describes a set of functional hypotheses proposed to explain the emergence of Rensch’s Rule in nature, the evidence supporting or undermining each of them, and potential research strategies that could increase the understanding of the mechanisms underlying allometric patterns of SSD like Rensch’s Rule in the light of these hypotheses.

The influence of these papers likely contributed to an exponential increase in the study of the allometry of male-biased SSD, now known as ‘Rensch’s Rule’ (Figure 2), and the study of SSD in general (Blanckenhorn, 2005). In the 30 years since the publication of the papers by Abouheif and Fairbairn, Rensch’s Rule has been tested in numerous taxa including amphibians, birds, crustaceans, fish, insects, primates and reptiles (e.g., Maly and Maly, 1999; Székely et al., 2004; Hansen et al., 2008; Serrano-Meneses et al., 2008; Steffen, 2009; Walker and McCormick, 2009; De Lisle and Rowe, 2013; Frydlová and Frynta, 2015). Although a common theme of the works studying the rule has been the test of the rule itself, many also shed light on the processes behind the pattern. For example, an important number of studies tested the importance of sexual selection in driving Rensch’s Rule, a mechanism suggested by Fairbairn and Abouheif (e.g., Dale et al., 2007; Serrano-Meneses et al., 2008). Others have tested the interaction between Rensch’s Rule and other macroecological rules, like Bergmann’s or Allen’s (e.g., Blanckenhorn et al., 2006; Lorrain-Soligon et al., 2023). Moreover, others have pushed forward our theoretical and conceptual understanding of the bases underlying Rensch’s Rule (e.g., De Lisle and Rowe, 2013; Adams et al., 2020; Meiri and Liang, 2021; Reyes-Puig et al., 2023), closing the gap between pattern and process.

**Figure 2.**
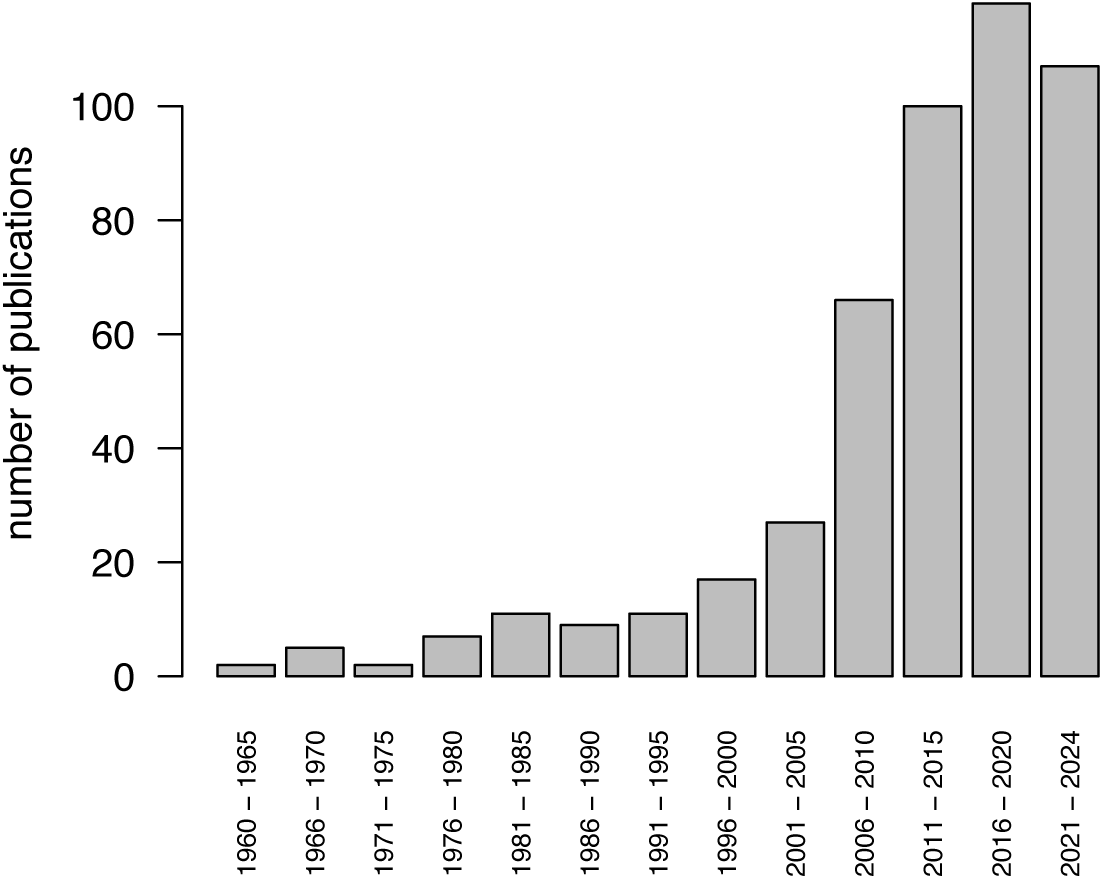
Number of academic documents published between 1960 and 2024 that cite Rensch 1950 and/or Rensch 1959 in the context of the allometry of sexual size dimorphism (i.e. Rensch’s Rule). The counting is based on academic documents (i.e., journal articles and theses) found through Google Scholar up to October 24th, 2024. Each bar corresponds to a five-year interval, except the last one.

Clearly, the popularization of Rensch’s Rule greatly contributed to our knowledge of the processes driving macroevolutionary patterns of phenotypic diversity, as well as to the general interest in the evolution of sexual size dimorphism. Its apparent ease to study and interpret has made of Rensch’s Rule one of the best known macroevolutionary patterns. However, as the papers by Abouheif and Fairbairn became citation classics for any work on Rensch’s Rule, some aspects of the original sources might have been overlooked. What exactly did Rensch discover and how does it relate to what we know today as Rensch’s Rule?

### II. Rensch’s original observations

#### Rensch 1950 – The dependence of relative sexual differences on body size

Two of Rensch’s works are considered the original sources of Rensch’s Rule and are cited by virtually all the studies testing the pattern. The first is a paper published in 1950 in German, titled *‘Die Abhängigkeit der relativen Sexualdifferenz von der Körpergrösse’* (translated as ‘*The dependence of relative sexual differences on body size*’). It was published in the *Bonner zoologische Beiträge*, nowadays known as the *Bonn Zoological Bulletin*.

Rensch started this paper citing the classic works of D’Arcy Thompson and Julian Huxley, explaining how different body parts follow different allometric trajectories throughout ontogeny, defining the body proportions observed at the different developmental stages of an organism. Rensch then suggested that these same processes of ontogenetic allometric scaling might be at work at the macroevolutionary level, shaping the morphological proportions of related species that differ in body size. To support this idea, he went on to provide a long list of patterns which describe how species of different sizes vary systematically in the proportions of different body parts (e.g., larger species of birds and mammals have relatively shorter ears, tails, and beaks; and relatively smaller brains, hearts, and eyes). This is important because it emphasizes the interest of Rensch in the proportions of particular body parts relative to the total size of the organism. In this context, Rensch introduced what he thought was a previously uncovered pattern:

> *“I would now like to add another [proportioning rule], which, however, will only be explained incompletely here, and which likely does not apply to all, but perhaps to the majority of animal groups: in species of the same family group the relative sexual difference is more substantial in the larger of these species than in the smaller ones.”*

This description resembles the modern definition of Rensch’s Rule. However, considering the context given in the first paragraphs, one could think that Rensch’s description refers mainly to dimorphism in particular body parts. Nonetheless, Rensch proceeds to exemplify his newly proposed ‘rule’ focusing on sexual size dimorphism (SSD):

> *“Let us first compare the most common sexual difference: the body size of the two sexes.”*

Here Rensch used examples from different groups of animals to test the rule. First, he showed that for each of several pairs of closely related German bird species the larger one tended to show higher male-biased size dimorphism. Nonetheless, he then showed how this trend is not observed among pairs of European mammal species, emphasizing the non-generality of this pattern.

Eventually, Rensch moved on to show how this allometric trend applies to different groups of carabid beetles. At this point, however, Rensch is not only comparing SSD between species of different sizes, but also how high levels of dimorphism in particular structures (e.g., the mandibles and tarsal segments of some beetles) appear only among the largest species. Overall, he considered SSD, dimorphism in individual body parts, and also the presence/absence of structures between sexes as different measures of dimorphism that, nonetheless, seemed to follow the same rule. He made this point very clear as he started closing his paper with a statement about the nature of his newly described rule:

> *“Different animal groups […] show a relatively greater sexual difference in larger species with regard to morphological sexual characteristics and body size.”*

Rensch was interested in the apparent macroevolutionary increase in sexual dimorphism with species size. But by using the term ‘sexual dimorphism’ he clearly did not discriminate between SSD and sexual differences in other traits (e.g., secondary sexual characters).

#### Rensch 1959 – Evolution Above the Species Level

*Evolution Above the Species Level* is the English translation of the second edition of the book *Neuere Probleme der Abstammungslehre*, written in German (Rensch, 1947, 1954), and is considered one of the most important works of the Modern Evolutionary Synthesis. Although the first German edition was published before the 1950 paper, the book only started including information on the allometry of dimorphism in its second edition (Rensch, 1954), on which the first English edition was based. Thus, here I focus on the text from the first English edition of *Evolution Above the Species Level*.

Rensch covers a lot of ground in this book, but it is still surprising that he wrote so little about the pattern he presented in detail to the world in 1950. Maybe more surprising is that Rensch seemed to ignore SSD altogether when describing his rule in this book, especially considering the modern definition of Rensch’s Rule. It is thus worth revisiting the few lines Rensch wrote about his newly described pattern in this book:

> *“Secondary sex characters of many animals usually represent structures of positively allometric growth because they usually become conspicuous during a more advanced period of ontogenetic development (due in mammals to the increasing influence of sex hormones). Thus, the rule is valid that in numerous animal groups the sexual dimorphism increases with body size (B. Rensch, 1950). This rule, however, applies only to subspecies of a species, to related species of a genus, or to related genera of a family.”*

So far, Rensch seems to be interested mainly in dimorphism in secondary sexual characters. No explicit mention of body size dimorphism so far. He also cites his 1950 paper. Let us look at the next lines:

> *“In species of birds in which the male is larger than the female, the relative sexual difference increases with body size. If, by way of exception, the females are larger than the males, as among many species of birds of prey, the opposite correlation applies, i.e. the greater sexual difference is found in the smaller species. This fact is probably due to the more negative allometry of sexual characters in the male.”*

Here, Rensch used the example of birds to describe how the dimorphism-size relationship changes depending on whether the focal species show male- or female-biased SSD. These ‘opposite correlations’ can actually be considered part of the same pattern (Meiri and Liang, 2021; Figure 1). Interestingly, these lines (except the last sentence) are quoted by Abouheif and Fairbairn (1997) as the original description of the modern size-focused Rensch’s Rule. They were not wrong, considering that Rensch included all types of dimorphism, including SSD, in his rule. However, Rensch probably referred to any kind of relative sexual difference, as shown in his 1950 paper. The last sentence of this fragment supports this idea, as it specifically refers to ‘the more negative allometry of sexual characters…’. The appearance of the original Rensch’s Rule in this book ends with a clear emphasis on secondary sexual traits as good examples for this ‘rule of sexual differences’.

> *“This rule of sexual differences is especially well demonstrated in some groups of beetles showing an excessive differentiation of the male antennae (as in some Lamellicornia; Figure 50) or of the tarsal sucking discs in the male (as in Dysticidae). In mammals, the anatomical structures used as weapons, such as antlers, horns, teeth, and the like, provide more examples of this kind. One need only compare large and small species of deer, pigs, baboons, and so forth.”*

Despite the disproportionate attention put on SSD by the modern Rensch’s Rule, it is clear that Rensch’s original pattern could be tested in, and was confirmed for, other kinds of dimorphisms. Rensch’s 1950 pioneering paper provides several examples of this, and *Evolution Above the Species Level* seems to focus on how dimorphism in secondary sexual traits follows the rule, without an explicit mention of SSD.

In the following decades, Rensch’s observations were regularly tested in different taxa, starting with birds (e.g., Amadon, 1959; Selander 1966; Earhart and Johnson 1970) before eventually including studies on mammals and reptiles (e.g., Schoener, 1969; Ralls, 1977; Gibbons and Lovich, 1990). Rensch’s pattern also influenced some landmark papers, including Lande’s model for the evolution of sexual dimorphism (Lande, 1980) and Cheverud’s, Leutenegger’s, and Clutton-Brock’s studies on sexual dimorphism and sexual selection on primates (Clutton-Brock et al., 1977; Leutenegger, 1978; Leutenegger and Cheverud, 1982, 1985; Clutton-Brock, 1985). However, as mentioned in the previous section, it was not until the 1990s that the study of Rensch’s Rule increased exponentially (Figure 2). Interestingly, prior to the 1990s most of the studies on Rensch’s Rule were already focusing only on the allometry of SSD, a pattern that would start to be amplified by the time Fairbairn and Abouheif’s papers were published.

The fact that Rensch was not only interested in the relationship between size and SSD, but in the allometry of any relative sexual difference has been noticed by a few others in recent decades (e.g., Cuervo and Møller, 2000; Seifan et al., 2009; Werner et al., 2016; Adams et al., 2020). Nonetheless, the overwhelming majority of Rensch’s Rule studies did and still do define the pattern exclusively as a relationship between size and male-biased SSD, which is only part of Rensch’s original observations. Are we missing something by focusing all our attention on SSD when studying and interpreting Rensch’s Rule patterns?

### III. Towards the future: testing Rensch’s Rule in traits other than size complements the study of sexual dimorphism

A strong emphasis on size dimorphism is not a bad thing in itself, as the study of Rensch’s Rule patterns in body size has provided us with important insights on the evolution of SSD in the context of climatic (e.g., Blanckenhorn et al., 2006; Tarr et al., 2019; Toyama et al., 2023), ecological (Lorrain-Soligon et al., 2023), and physiological constraints (Stillwell et al., 2010). Nonetheless, the exploration of allometric patterns of relative dimorphism in individual body parts (i.e., testing Rensch’s Rule in morphological traits other than size) can improve our understanding of the processes underlying the evolution of sexual dimorphism.

To illustrate this, we can focus on sexual selection hypotheses for the emergence of Rensch’s Rule, which are frequently invoked when trying to explain allometric patterns of SSD (Webster, 1992; Andersson, 1994; Fairbairn, 1997; De Lisle and Rowe, 2013). Under sexual selection hypotheses, large male sizes are selected for in species that exhibit intrasexual mate competition, as large sizes frequently provide males advantages in competitive performance against other conspecific males (e.g., Clutton-Brock et al., 1977; Andersson, 1994; Haley et al., 1994). In this context, a positive relationship between male-biased SSD and species size (i.e. a Rensch’s Rule pattern) can arise because (a) species under stronger sexual selection become larger due to direct selection on male body size (and also due to the effects of correlated selection and/or genetic correlation on female body size), or because (b) the effects of sexual selection on male body size are stronger on larger species (Webster, 1992; Andersson, 1994). Regardless, it is improbable for male body size in a given species to be defined solely by the intensity of sexual selection. In fact, body size is most likely the result of a balance between several selection pressures acting simultaneously (Peters, 1983; Schmidt-Nielsen, 1984; Reiss, 1989). This implies that, under certain scenarios, the evolution of larger male sizes could be constrained such that a Rensch’s Rule pattern might be weak or even absent despite the presence of intrasexual selection in males. However, in such cases sexual selection might still drive the evolution of individual body parts that influence competitive performance, (e.g., the proportional dimensions of secondary sexual traits or weapons) even if body size is constrained. In those scenarios, testing Rensch’s Rule in those traits might still allow us to detect the signature of sexual selection. I will next show some empirical examples in which testing for Rensch’s original ‘rule of sexual differences’ in size and other traits can inform us of selection processes that would otherwise be overlooked.

#### Harvestmen and weevils: when body size and weapon size tell different stories

Species of harvestmen from the subfamily Mitobatinae (part of Opiliones, an order of arachnids) show female-biased SSD. However, males have a disproportionately elongated fourth pair of legs which they use to compete in signaling contests with other males (Machado et al., 2021). In this group of species, body size does not follow Rensch’s Rule (Figure 3A). However, the length of the fourth pair of legs does, as expected based on a sexual selection hypothesis for its exaggeration (Figure 3E). Similarly, species of weevils from the Brentinae subfamily (part of Brentidae, a family of coleopterans) show low levels of SSD on average, but males have much larger mandibles than females, which they use to fight other males (Painting et al., 2024). Like in harvestmen, the body size of these weevils does not follow Rensch’s Rule (Figure 3B), but the size of their mandibles does (Figure 3F). In both cases this means that the evolution of relatively larger secondary sexual traits in males, but not of disproportionate larger male sizes, is associated with the evolution of larger body sizes across species.

**Figure 3.**
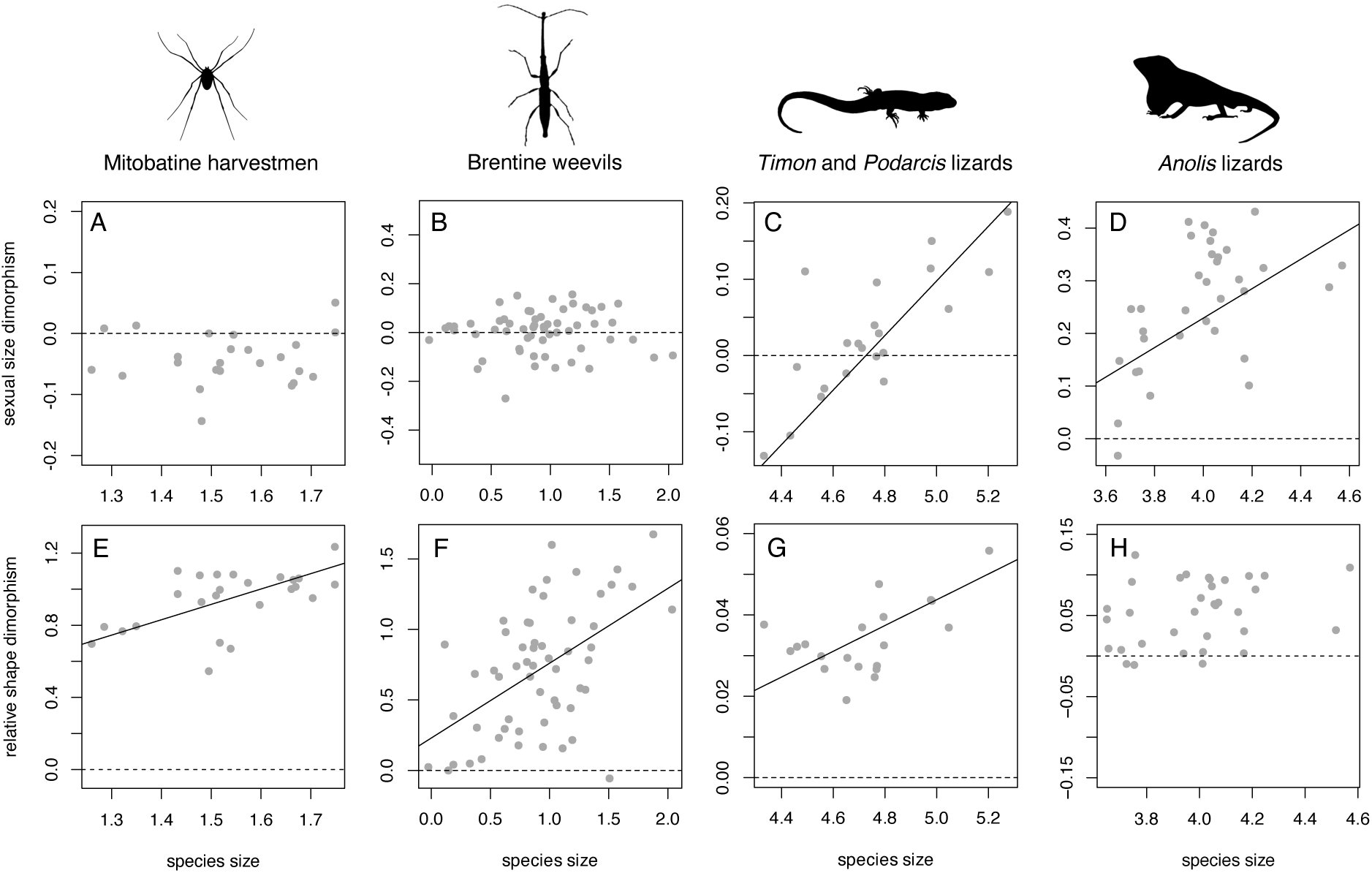
Allometric patterns of dimorphism in size and shape can differ. Mitobatine harvestmen (Opiliones) do not follow Rensch’s Rule in body size (A) but the length of their fourth pair of legs, which males use in signaling contests, do (E). Similarly, the body size of brentine weevils does not follow Rensch’s Rule (B), but the width of their mandibles, which males use to fight each other, does (F). Green lizards (*Timon* and *Podarcis* species) show Rensch’s Rule in both size (C) and head size (G), suggesting that males obtain relative performance enhancements from larger bodies and relatively larger heads. Insular anoles follow Rensch’s Rule in body size (D) but head size does not (H), suggesting that fighting performance in males is mainly driven by body size. Dashed lines indicate a value of dimorphism = 0. Positive values in the y-axes correspond to male-biased dimorphism. Solid lines represent significant fitted phylogenetic regressions. Data reanalyzed from Machado et al. 2021 (harvestmen), Painting et al. 2024 (weevils), Adams et al. 2020 (green lizards), and Toyama et al. 2024 (anoles).

Several mechanisms can be proposed for the evolution of Rensch’s Rule patterns in sexually selected traits (e.g. Webster, 1992; Andersson, 1994), some of which would have to be erroneously discarded if we were to base our inferences only on the patterns followed by body size in organisms like harvestmen or weevils. Instead, we can now combine the information provided by body size and other traits to ask further questions about these systems. Why are relatively long legs and relatively large mandibles more likely to evolve in males of the largest species of harvestmen and weevils, but relatively large male sizes are not? What are the roles of natural and sexual selection in promoting and constraining the evolution of dimorphism in size and shape?

#### Lizards: similar systems can show different patterns

Lizards represent ideal systems in which to study the evolution of sexual dimorphism under sexual selection. Male lizards from territorial species often compete with other males for access to reproductive opportunities and the outcomes of these interactions can often be predicted based on the sizes of the individuals involved (Jenssen and Nunez, 1998; Sacchi et al., 2009). Since male lizards often fight using their jaws to bite each other (Huyghe et al., 2005; Losos, 2009), larger individuals have an advantage due to their stronger absolute bite forces. Green lizards (a group that includes *Lacerta* and *Podarcis*, Adams et al., 2020) represent a typical group of territorial lizards in which male-male antagonistic interactions are common. As expected, males are often larger than females in species of this group and Rensch’s Rule in body size is followed (Figure 3C). Moreover, head size in these lizards also follows a Rensch’s Rule pattern (i.e., the relative sexual difference in the robustness of the head increases with body size across species, Figure 3G). This suggests that sexual differences in fighting performance (i.e., biting) are likely more pronounced in larger species due to the evolution of both proportionally larger bodies and proportionally larger heads in males from those species. In other words, the influence of sexual selection is expressed and can be detected in both body size and head proportions.

*Anolis* lizards share some characteristics with green lizards: most anole species show male-biased SSD, many have been shown to be territorial, and males engage in combat for access to mating opportunities also using their jaws to bite each other (Johnson et al., 2010; Losos, 2009). As expected, body sizes in anoles also follow Rensch’s Rule (Figure 3D – for clarity, only species from insular clades are shown, but the pattern is also followed by mainland species, Toyama et al., 2024). However, in contrast to *Podarcis* and *Lacerta* species, head size does not follow a Rensch’s Rule pattern, even though heads are consistently larger in males across species (Figure 3H). These patterns indicate that sexual differences in biting performance should be more pronounced in larger species mainly due to the evolution of larger male body sizes, but not due to the evolution of relatively larger male heads in those large species.

The reasons why different lizard radiations (or radiations of other closely-related taxa) might show different patterns of allometry in dimorphism will need to be addressed in future studies, but more integrative tests of Rensch’s Rule allow new hypotheses to be proposed. For example, the arboreal ecology of most anole species might represent a constraint for the evolution of exaggerated head dimensions (which define biting performance, Herrel et al., 2001; De Meyer et al., 2019), as large heads can quickly become detrimental for vertical locomotion and habitat use (Vanhooydonck and Van Damme, 1999; Revell et al., 2007). This hypothetical constraint would thus be hindering the evolution of additional performance enhancements in males in this group of arboreal lizards.

These examples suggest that, like body size, secondary sexual traits can also be constrained by natural selection (Tidière et al., 2017; Painting et al., 2024); and how, again, exploring the allometry of dimorphism in more than just body size can shed light on the processes underlying the evolution of sexual differences.

#### A final cautionary note

The study of Rensch’s Rule patterns on traits other than body size will likely improve our understanding of selection processes involved in the evolution of dimorphism. However, some caution needs to be exercised if this kind of studies are to increase, especially regarding the methods to be used and their interpretation. The log-log regression method popularized by Abouheif and Fairbairn (Abouheif and Fairbairn, 1997; Fairbairn, 1997) is useful when testing Rensch’s Rule in body size because it can simultaneously illustrate species size and SSD (Figure 1B). However, using this same method on other traits is not analogous to using it with body size and should not be considered a test of Rensch’s Rule.

I will illustrate this with the *Anolis* head size example. As shown before, the average difference in relative head size between males and females does not change with species size in anoles (Figure 3H) indicating that head size does not follow Rensch’s Rule. However, if we were to test Rensch’s Rule looking at the relationship between log-transformed male head size and log-transformed female head size, a slope > 1 would seem to confirm the rule (Figure 4A). Although intuitive, this analysis does not test the presence of an allometric pattern because it does not include a direct measure of body size. Even if head size is positively associated with body size, the variability around a head size ∼ body size relationship might result in discordance between both analyses because some species might have larger or smaller head sizes for their body size. Thus, a log-log test in this case is not accurate regarding the allometry of relative head size dimorphism, but explores whether species with absolutely larger heads (but not necessarily larger bodies) are the most dimorphic in head size.

**Figure 4.**
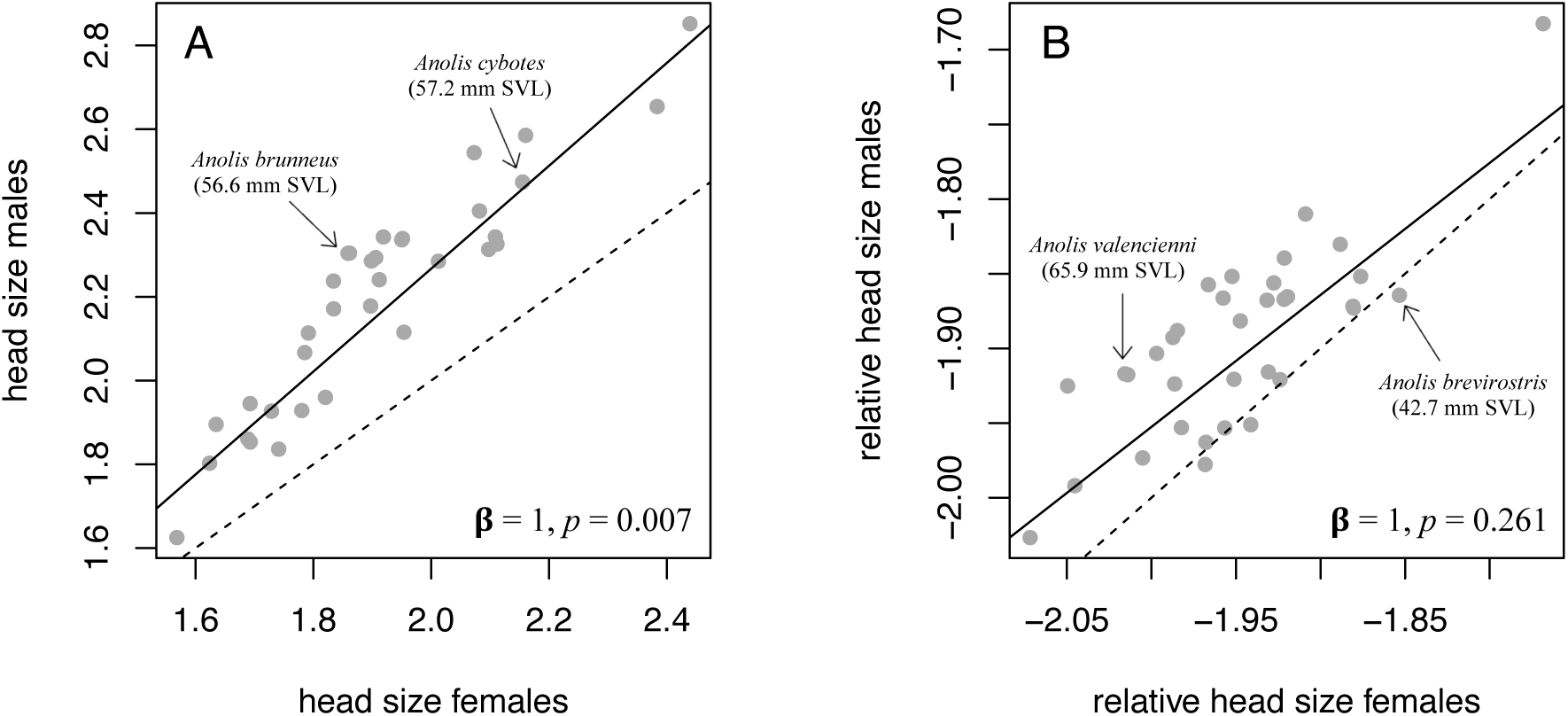
Regressions between male and female body parts are not testing Rensch’s Rule. (A) A slope > 1 in a regression of log-transformed male on log-transformed female head size indicates that, in species with larger absolute head sizes, males have larger heads than females. This does not accurately represent a test of Rensch’s Rule because a measure of body size is not explicitly included in the analysis, even if head size is positively associated to body size. Notice how two anole species (arrows in figure) can differ in average head size while having similar average body sizes. An analysis that directly tests the association between relative head size dimorphism and species size shows no evidence of Rensch’s Rule (Figure 3H). (B) A regression between male and female relative head size is even farther from an appropriate test of Rensch’s Rule because body size is removed from both variables. Notice how anole species with relatively larger head sizes can have smaller body sizes than species with relatively smaller head sizes (arrows in figure). In this analysis we can only conclude that the level of relative (i.e. size-corrected) head size dimorphism is independent of the relative (i.e. size-corrected) head size of the species. In both panels, a null hypothesis in which the slope of the relationship is 1 was tested with *F* statistics (results shown in each panel). Dashed lines represent 1:1 relationships, solid lines represent fitted phylogenetic linear models between male and female traits.

Similarly, we might apply the log-log method to the relative size of the head (i.e., head size corrected for body size). In this case, the slope of the regression between male and female relative head size is not different from 1 (Figure 4B). However, this analysis is even farther from a Rensch’s Rule test than the previous one because size is completely absent from the variables involved. For example, in Figure 4B the species near the upper-right corner of the plot are not the ones with the largest heads or bodies, but the ones with the most disproportionally large heads relative to their body size. In this case, we cannot infer anything about the allometry of dimorphism. This analysis, however, can be useful in answering different questions. For example, this result indicates that dimorphism in relative head size is independent of relative species head size. In other words, the species with proportionally larger heads are not more sexually dimorphic in relative head proportions than species with proportionally smaller heads (see Juarez and Adams 2022 for an example of this kind of analysis).

Rensch’s Rule, both as originally conceived by Rensch and as understood by most researchers in recent times, is a pattern of allometry. It should describe the relationship between a variable (in this case, relative sexual dimorphism) and a measure of body size. If Rensch’s Rule is considered a pattern of interspecific allometry, is vital to include a direct measure of body size when testing it.

## Conclusion

Since Rensch first defined his ‘rule of sexual differences’, an overwhelming majority of subsequent studies focused on how this rule applied to sexual dimorphism in body size. More than seven decades later, this bias has resulted in ‘Rensch’s Rule’ being almost exclusively defined as an allometric pattern of sexual size dimorphism (SSD). Interestingly, Rensch’s original observations were not centered on SSD and, if anything, Rensch found stronger empirical support for his canonical pattern from individual body parts. The study of SSD in the context of Rensch’s Rule is, of course, not bad in itself, as I mentioned above. However, empirical data made evident that testing Rensch’s Rule on other traits in addition to body size can increase our understanding of the processes and mechanisms driving the evolution of sexual dimorphism, or at least foster the exploration of new hypotheses.

What should we do about the meaning of the term ‘Rensch’s Rule’? To be historically accurate, Rensch’s Rule should not be assumed to involve dimorphism in body size, as the pattern can also be tested on the relative sexual differences observed in any particular body part. In this text I have used expressions like ‘Rensch’s Rule in body size’ or ‘head size does not follow a Rensch’s Rule pattern’. My intention was to separate the pattern from the trait, as Rensch did when originally describing the pattern. Nonetheless, I’m aware that it might be hard to redefine a concept that has been consistently used for such a long time. It will be more important for researchers to become aware of the possibility of looking at ‘Rensch’s Rule-like’ patterns in traits other than body size, and to do so with the appropriate methods, as I briefly exemplified in the previous section.

The study of Rensch’s Rule in individual body parts has yet to take off, and the decades of theoretical and empirical advances obtained from its application to a single trait should give us a perspective of the countless new puzzles that await to be solved.

## Acknowledgements

I want to thank Luke Mahler, Miriam Ahmad-Gawel, Alexander Tinius, Jill Sanderson, Gavia Lertzman-Lepofsky, Rowan French, and Zifang Xiong for comments, suggestions, and corrections in an earlier version of this manuscript. I am especially grateful to Alexander Tinius for his help with translating text from German.

